# Effects of Weather and Season on Human Brain Volume

**DOI:** 10.1101/2020.07.07.191239

**Authors:** Gregory A. Book, Shashwath A. Meda, Ronald Janssen, Alecia D. Dager, Andrew Poppe, Michael C. Stevens, Michal Assaf, David Glahn, Godfrey D. Pearlson

## Abstract

We present an exploratory cross-sectional analysis of the effect of season and weather on Freesurfer-derived brain volumes from a sample of 3,279 healthy individuals collected on two MRI scanners in Hartford, CT, USA over a 15 year period. Weather and seasonal effects were analyzed using a single linear regression model with age, sex, motion, scan sequence, time-of-day, month of the year, and the deviation from average barometric pressure, air temperature, and humidity, as covariates. FDR correction for multiple comparisons was applied to groups of non-overlapping ROIs. Significant negative relationships were found between the left- and right-cerebellum cortex and pressure (t = −2.25, p = 0.049; t = −2.771, p = 0.017). Significant positive relationships were found between left- and right-cerebellum cortex and white matter between the comparisons of January/June and January/September. Significant negative relationships were found between several subcortical ROIs for the summer months compared to January. An opposing effect was observed between the supra- and infra-tentorium, with opposite effect directions in winter and summer. Cohen’s d effect sizes from monthly comparisons were similar to those reported in recent psychiatric big-data publications, raising the possibility that seasonal changes and weather may be confounds in large cohort studies. Additionally, changes in brain volume due to natural environmental variation have not been reported before and may have implications for weather-related and seasonal ailments.

## Introduction

Studies testing the effects of weather and season on the human body have found relationships between these environmental factors and incidence or severity of disease. Sales of headache medicines increase when barometric pressure decreases, and spontaneous delivery rates increase when barometric pressure drops [1, 2]. Environmental effects on specific diseases have been studied, including multiple sclerosis (MS), schizophrenia, and Alzheimer’s. A significant relationship exists between winter conditions and a higher incidence of onset or recurrence of multiple sclerosis [3]. A similar relationship exists between MS relapse rates and latitude, with rates increasing further from the equator [4]. An association has been observed between season and first-break schizophrenia and psychosis, with a stronger effect in males [5, 6]. Seasonal rhythms in gene expression have been found to be interrupted by Alzheimer’s disease [7]. Season and weather associations with symptoms of particular disorders have been investigated from a public health perspective, but the underlying biological response to environmental factors has not been as extensively studied. A study showed an association between hippocampal volume and photoperiod [8]. One study examined the effect of time-of-day on a longitudinal sample of 755 MS and 834 Alzheimer’s patients [9], while a second study examined a controlled sample of 19 healthy young adults [10]. Both studies found that total brain volume decreases throughout the day. Studies of cognition have found a seasonal periodicity associated with task performance [11, 12].

Human response to environmental rhythms may be similar to that of other animals, as animals alter their physiology to adapt to changing seasonal energy needs. Animal studies have found seasonal structural changes to the hippocampus, total brain volume, and cranium size in mammal, amphibian, and avian species [13–17]. A study of seasonal changes in the brain volume of the common shrew found the cerebellum increasing in volume by 8.0% from summer to winter and the rest of the brain decreasing in volume by 11.5% [13]. The tentorium appeared to act as a divider between effect directions in this study, and much larger effects were observed in males.

Change in daylight is a significant factor in seasonal studies, but few studies have taken into account weather conditions, and no studies have examined the effects of weather on brain volume. Weather is often described as temperature, precipitation, and wind speed, but the most significant driver of weather is barometric pressure. Air moves from areas of high pressure to low bringing with it wind, and changes in temperature and precipitation. Unlike temperature and humidity which are well-controlled in MRI scanning environments, pressure is ubiquitous and thus a good weather variable to explore. A phenomenon similar to changes in barometric pressure that has been studied is the effect of high-altitude exposure on brain volume. High-altitude (HA) exposure has been studied in humans and measurements of brain volume have been conducted. A three month HA exposure caused an increase in brain volume in one study [18]. At sea-level pressure, but in zero-gravity, cosmonauts were found to experience brain volume changes after 189 days in space [19].

Exploring seasonal and weather changes in brain volume is best analyzed using a very large dataset. Using a sample of healthy control subjects collected at the Olin Neuropsychiatry Research Center, located in Hartford, CT USA over a 15 year period, we explored the effects of environmental factors of season and weather on brain volume. Hartford is an ideal location to test weather and seasonal effects because it is near sea-level, experiences four distinct seasons, and a wide range of weather conditions. Because weather is highly correlated with the season, we attempt to separate the effects of weather and season. We additionally compare the effect sizes found in this study to those found in large-scale neuroimaging studies, and attempted to replicate previous findings of a diurnal effect on brain volume and a change in hippocampal volume based on time of year.

## Materials and Methods

### Imaging Data Collection & Processing

Imaging data was gathered retrospectively from approximately 12,600 structural T1-weighted MRI scans collected between August 2003 and October 2018 at the Olin Neuropsychiatry Research Center, Institute of Living, in Hartford, CT USA. Subjects who received MRI scans were recruited into individual neuropsychiatric studies. Those subjects received a complete description of the studies in which they participated, and written informed consent was obtained prior to scanning. Scans were performed on a Siemens Allegra 3T head-only MRI and a Siemens Skyra 3T MRI scanner (Siemens Medical Solutions, Malvern PA). Six structural T1 MRI pulse sequences were used between the two MRI scanners (table 4). Images were analyzed automatically using Freesurfer 6.0 [20] and the `recon-all` command with `-all` and `-notal-check` options. Computational analysis was performed using an instance of the Neuroinformatics Database (NiDB) [21] and took 195,000hrs (22.5 years) of CPU time to complete on a 300-core Linux cluster. Subcortical and summary regions of interest were extracted using the default automatic subcortical segmentation (aseg) atlas [22]. Summary ROIs used for analysis included BrainStem, SubCortGrayVol, CortexVol, and CerebralWhiteMatterVol. Lateral ROIs used for analysis included left- and right- amygdala, caudate, cerebellum cortex, cerebellum white matter, hippocampus, pallidum, putamen, thalamus, cerebral white matter, and cerebral cortex. All ROIs were corrected for estimated total intracranial volume (eTIV), to remove effects of head volume.

Recent publications indicate that head motion is associated with a decrease in Freesurfer volumes. To account for possible subject motion, a motion metric was calculated for each subject’s dataset using the methods described in the paper by Reuter et al [23]. Motion was estimated from any fMRI timeseries collected in the same imaging study as the T1 with at least 100 time points collected. Timeseries data may have been a task or resting state scan. These motion estimates were calculated by performing rigid realignment using FSL’s MCFLIRT tool [24]. The derivative of the resulting motion correction was calculated, giving a displacement value in mm between adjacent time points, which ignores the effect of slow physical motion in the scanner. Root mean square (RMS) of the maximum displacement in the x, y, and z-directions were calculated, with the largest value used as a ‘motion’ variable for later statistical analysis.

After processing of the imaging data through Freesurfer and FSL, cleaning and quality control was performed. Exclusion criteria included: individuals with invalid ages, invalid/unknown sex, incidental findings (tumor, aneurism, AVM, etc), history of traumatic brain injury, enrollment in pre-surgical mapping studies, and incomplete Freesurfer analyses and/or fMRI data. Arbitrary cutoffs, determined from visual inspection of the data, were used to exclude analyses with outlying results; datasets with a BrainSegVol-to-eTIV (estimated total intracranial volume) ratio of greater than 1.05 or less than 0.6 were excluded, as well as eTIV’s less than 900,000 mm^3^. Due to the size of the remaining sample, hand-editing of Freesurfer segmented surfaces was not performed. However, rendered images of pial surface maps were reviewed and incorrectly segmented subjects were excluded. Thumbnails of raw T1 data were also examined and subjects with visible artifacts (usually motion related) were excluded. For subjects with more than one scan, only the most recent MRI scan was included to attempt to balance the sample away from a younger average age. After all quality control and data cleaning, 6,139 subjects remained.

Imaging data was pooled from over 150 separate research projects that primarily studied psychiatric disorders – each with different enrollment criteria, different definitions of *healthy*, *control*, and *patient*, and differing levels of detail for diagnoses. Some individuals received a full structured clinical interview (SCID) to determine DSM diagnosis, but most participants did not undergo formal psychiatric diagnostic interview. Many individuals did not receive a diagnosis but were enrolled in projects that solely enrolled “healthy” participants. Subjects with diagnosis labels of schizophrenia, bipolar, psychosis, major depression, Alzheimer’s, traumatic brain injury, and autism were excluded. Subjects who were not explicitly labeled as “healthy” but were enrolled in projects which also enrolled those diagnoses were excluded from analysis. 4,039 subjects explicitly labeled, or implicitly defined as, “healthy controls”, remained. The remaining sample included subjects ranging in age from 9-93 years. To remove possible pediatric effects subjects younger than 18 were excluded, and to balance the mean age between seasons, subjects older than 65 were excluded, leaving 3,279 healthy individuals for analysis.

### MRI Quality Control Data

MRI quality control (QC) data was collected semi-regularly over the course of the analysis period. QC MRI scans were collected on the Allegra MRI using an MPRAGE (multiplanar rapid acquisition gradient echo) pulse sequence (256×240×260 voxels, 1.3×1×1mm voxel size, 2300ms TR, 2.91ms TE, 9° flip angle) on an ADNI phantom (The Phantom Laboratory, https://www.phantomlab.com/magphan-adni). QC scans were collected on the Skyra MRI using an MPRAGE sequence (176×240×256 voxels, 1.1×1.1×1.2mm voxel size, 2300ms TR, 2.95ms TE, 9° flip angle) on an ACR small phantom (Newmatic Medical, Caledonia, MI). Signal-to-noise ratio (SNR) was calculated by dividing the signal (mean intensity of non-noise areas) by the noise (mean intensity of the corners of the image volume).

### Environmental Data

Weather data was obtained using the National Oceanic and Atmospheric Administration’s (NOAA) Local Climatological Data (LCD) search tool for the period of August 4, 2003 to October 30, 2018, from Bradley International Airport, which is the closest weather station with contiguous data for the time period (https://www.ncdc.noaa.gov/cdo-web/datatools/lcd). Bradley Airport is located 12 miles (20km) from the MRI collection site and has an elevation of 170ft (51.8m). The Olin Center’s elevation is approximately 110ft (33.5m). The LCD dataset contained hourly weather variables used in the analysis: Dry Bulb Temp (temperature in C), Relative Humidity (humidity in %), Station Pressure (barometric pressure in inHg). The nearest hourly measurement to the start time of the T1 scan was used in analysis.

Köppen climate classification identifies Hartford, CT, USA as a humid continental climate (Dfa) characterized by hot summer, cold winter, and well distributed year-round precipitation, with four distinct seasons [25]. For simplicity, astronomical season was defined as starting on the 21^st^ day of March, June, September, and December, so that days of the year 80-171 were labeled spring, days 172-263 labeled summer, days 264-354 labeled fall, and all other days labeled winter. Scan time-of-day was obtained from the DICOM header for the T1 series. Because time-of-year, temperature, and humidity are highly correlated, we attempted to separate the effects of time of year and weather by using the deviation of weather variables from monthly mean. Mean monthly temperature, pressure, and humidity were calculated over the 15-year period, from which the deviation from the monthly averages of weather at individual scan time-points was calculated. This deviation from monthly mean was then used in analysis and referred to as *pressure*, *temperature*, and *humidity*. This method distinguishes the effects of time-of-year from the effects of the departure from normal weather conditions; ie, is an effect of temperature because temperature is hottest in July or because of warmer than average temperatures on any given day of the year.

### Statistical Analysis

We wanted to determine if effects seen were due to weather or time-of-year, so a linear model was used where each FreeSurfer ROI served as a dependent measure, and age, motion, sex, time-of-day, and deviation from pressure/temperature/humidity were independent continuous variables, and scan sequence and month were categorical variables. ROIs were selected because they were whole-brain or summary regions (total gray matter, cortex volume, etc) or defined structures (amygdala, putamen, etc) that have been implicated in the limited prior literature. ICV may be change with age, but was not included in the model because of its strong correlation with age. Because of previous evidence of sex differences in brain volume, similar analyses were performed for only males and only females. Analyses were performed using the R statistical software package (http://r-project.org) and significant results with *p* < 0.05 were noted, using FDR correction for multiple comparisons across non-overlapping groups of ROIs. Additional post-hoc t-tests were performed for each ROI for a month-month comparison. Uncorrected p-values less than 0.05 were noted. Percent difference in volume from mean, and the Cohen’s *d* effect size, of the factor of interest, between months were calculated. For graphical purposes, monthly percent different from annual mean were calculated for each ROI.

Body-mass index data was only available on 517 of the 3279 subjects included in the main analysis. A separate analysis of that subset, using BMI as a covariate was performed, and the results included in supplement tables 2 and 3.

## Results

### Subjects

Subjects ranged in age from 18 to 65, with a mean age of 32.4 (+/− 13.5) years; 1,779 female (33.4 +/− 14.2 years), and 1,500 male (31.3 +/− 12.6 years). Pairwise t-tests by month, FDR corrected for multiple comparisons, showed no significant differences in scan sequence or motion. Significant differences were found in one month-month comparison for sex (supplement table 1b), and eight month-month comparisons for age (supplement table 1a), and no significant differences for motion or scantype by month. MRI quality control data did not indicate an association between phantom SNR and time of year.

### Weather

Weather data was available within +/− two hours for 91.4% of the scans in the dataset. For the remaining datasets, the nearest weather measurements within six hours were used. Minimum and maximum measurements of pressure, temperature, and humidity during MRI scanning ranged from - 16.1C to 38.3C, 10% to 100%, 28.82 inHg to 30.51 inHg respectively.

### Weather and seasonal effects

Pressure was negatively associated with supra-tentorial and caudate volumes, while cerebellum cortex and white matter volumes were positively associated with pressure (table 1). Temperature and humidity were not associated with changes in any brain regions. Several ROIs showed significant associations with January-June and January-August comparisons (table 1). Seasonal percent-different-from-mean in males and females were different (figure 1), particularly that cerebellum volume peaks in females in June, and peaks in males in September (figure 1A). Effects of pressure were only found in females, and only in the supra-tentorial, left/right cerebellum, and right cerebellum white matter. More subcortical ROIs were significantly different than January for the month of July in males and August in females (tables 2, 3). *Post-hoc* uncorrected t-tests by month showed subcortical gray matter volume decreased between January and August (*p* = .003, Cohen’s d = −.228) and increased between August and December (*p* = 0.013, Cohen’s d = 0.203). Left- and right- cerebellum cortex increased in volume between January and June (*p* = .003, Cohen’s d = .221; *p* = 0.011, Cohen’s d = 0.202) and decreased between July and December (*p* < .001, Cohen’s d = −0.262; *p* = 0.007, Cohen’s d = −0.211) decreased during the same period. The tentorium acted as a divider between effect direction, with changes from summer to winter months being positive for supra-tentorial ROIs and negative for infra-tentorial ROIs (table 1).

**Figure 1.**
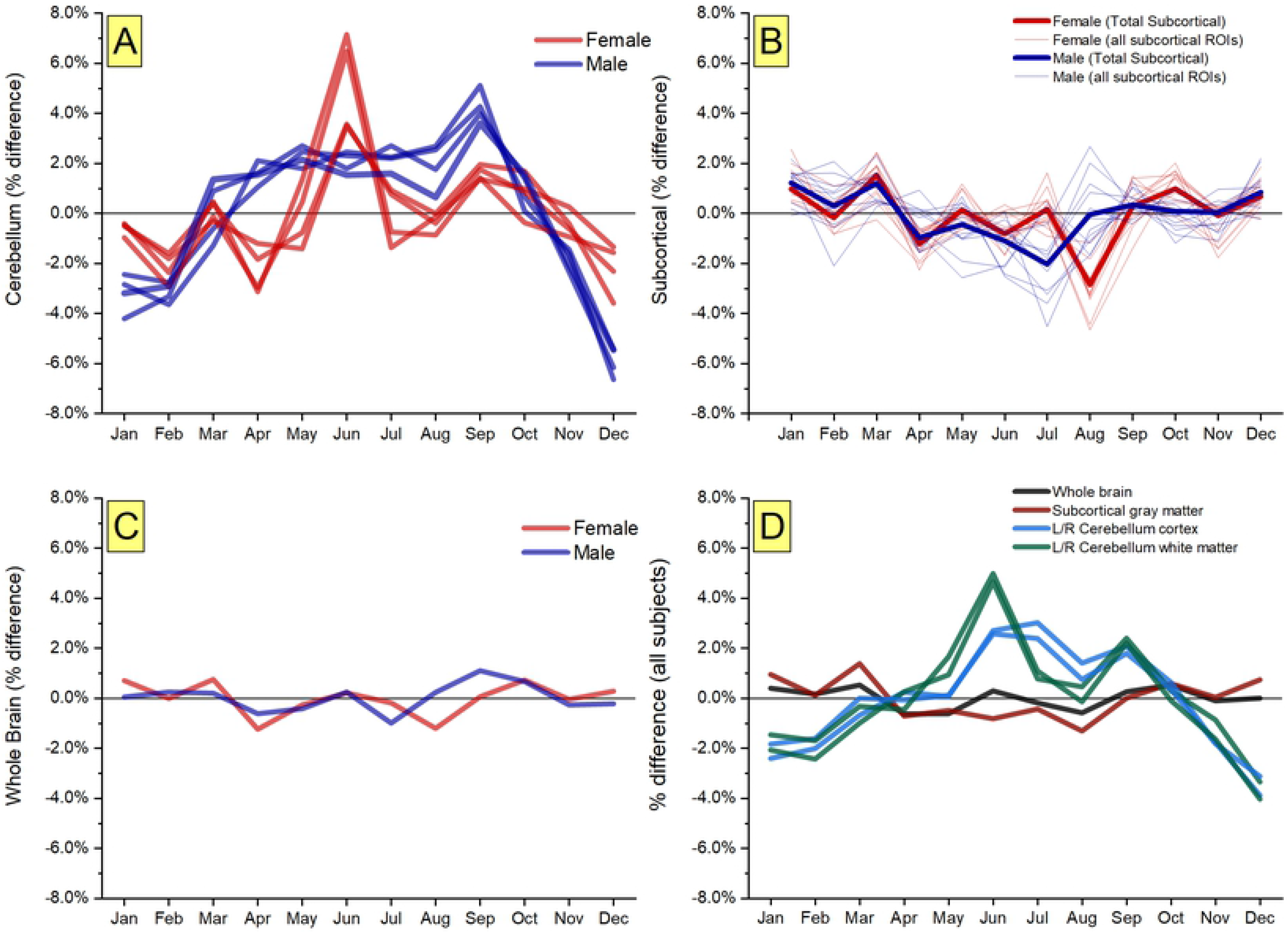
Monthly volumes – difference from yearly mean [A] Cerebellum cortex and cerebellum white matter in males and females. [B] Total subcortical gray matter volume for male and female represented by thick lines, and individual subcortical ROI volumes represented by thin lines. [C] Whole brain volume for males and females. [D] Summary volumes for all subjects.

## Discussion

Season and weather have a known and appreciable effect on the human body, and we have found evidence of previously unmeasured changes in brain volumes. We were unable to replicate a significant time-of-day effect on any brain volume ROI but were able to replicate a seasonal effect on hippocampal volume, though only in females. Time-of-day changes reported in other studies were attributed to hydration status, and we hypothesized that extremes of weather such as a hot dry day, or cool humid day may be reflected in brain volume. No association was observed between time-of-day and brain volume. When controlling for other factors, changes in humidity and temperature from normal had no effect on brain volumes. Results from other studies found mixed results on whether hydration status significantly changes the brain volumes measured from MRI images [26, 27].

### Environmental Factors as Confounds

There is increasing interest in aggregating large samples of psychiatric patients to look for evidence that specific psychiatric diagnoses might have different brain structure than non-patient samples [28]. Brain volume changes observed in this study reveal a possible confound in this approach to big data analysis. Effect sizes observed from the seasonal changes in brain volume in this study were in some cases larger than the effect sizes of patient/control comparisons in recent big-data analyses. This potentially represents a considerable confound when drawing conclusions about patient/control population if these other sources of variation are not controlled. Changes in environmental temperature and barometric pressure are known to affect blood pressure and oxygen saturation and are considered confounds to accurate vital sign measurement in clinical environments [29, 30]. Such confounds from barometric pressure and season may also exist in neuroimaging studies.

A comparison of effect sizes can be made between this study and those of previously published large-scale studies, using the ENIGMA consortium as example. ENIGMA has published several large-scale studies comparing Freesurfer derived ROI volumes between controls and patients for various disorders. An analysis of 2,028 schizophrenia patients and 2,540 controls found significant differences in hippocampus, amygdala, thalamus, and lateral ventricles [31]. The effect size of the differences in thalamus volume between populations in the ENIGMA analysis was 0.31 (2.74% difference), compared to a Cohen’s d effect size between March and August in the left- and right- thalamus in the Olin sample of 0.213 (2.98%) and 0.216 (2.94%). A comparison of 1,728 major depressive disorder (MDD) patients and 7,199 controls showed a significant difference between populations in the hippocampus (1.25%, Cohen’s *d* = 0.144) [32]. Differences were described in the amygdala, hippocampus, and thalamus in another paper comparing 1,026 epileptics and 1,727 controls [33]. A comparison of 2,140 substance users and 1,100 controls found differences in the amygdala, hippocampus, putamen, thalamus [34]. Results from these four papers are listed adjacent to the largest effects found in this study (table 5).

When performing large sample analyses, many unknown factors may influence results. The current results suggest that data collection should be uniform across season, but also that it should be standard practice to statistically model for variation due to season in geographical areas where seasons are distinct and widely variable. It is likely important to include barometric pressure as a statistical covariate when using data gathered from climatologically diverse data collection sites. It is entirely possible that a case-control research study recruits most of the patients at the start of the project in the winter and fills in the controls the following summer. Scanning more subjects of one group in a season might represent the effect seen in a case-control analysis, especially as the effects observed in these analyses are already somewhat small.

### Biological Significance

Many health effects and diseases are associated with season or weather. Approximately 5% of the US population experiences seasonal affective disorder in a given year [35]. Headache may have a weather-related trigger as indicated by significantly higher sales of over-the-counter headache medications when barometric pressure dropped the previous day [36]. Studies of the influence of weather on migraine have shown mixed results [1, 37, 38]. Though not neurologically related, drops in barometric pressure cause an increased risk of spontaneous cephalic delivery [2]. Our findings indicate that changes in barometric pressure have a larger effect on the brain volume of females than males, so barometric pressure changes may affect females in multiple ways.

A surprising finding from this study was that supra-tentorial regions gain volume when ‘bad’ weather approaches - either when barometric pressure drops or when winter is coming - but cerebellum and brain stem volumes change in the opposite direction. While these countervailing effects on different parts of the brain defy easy explanation, they are not without precedent in mammals. Such seasonal brain volume changes are similar to those of the common shrew, with the tentorium acting as a divider between effect directions and with males having larger changes than females. Volume change directions in the shrew are opposite that humans based on time-of-year, however that may be dependent on the average one-and-a-half year lifespan of the shrew. Blood to infra- and supra-tentorial regions are supplied by different vasculature, which may be responsible for opposite changes in volumes. A strong seasonality effect raises a possible explanation of changing vitamin-D levels. Previous studies have found negative association between vitamin-D levels and intracranial volume, and vitamin-D and season. Subjects with lower levels of vitamin-D showed larger intracranial and white-matter volumes [39], and lower levels of vitamin-D are found in winter [40]. Though this hypothesis is speculative, it is testable in subjects who are prescribed light exposure during winter months for various conditions.

A possible explanation of brain volume changes is from a change in blood flow as previous studies have found a seasonal effect on ambulatory blood pressure [41, 42]. Blood flow associated with barometric pressure may also offer an explanation. The decrease in barometric pressure associated with an increase in infra-tentorial volumes found in this study may be explained by a vascular response to available oxygen levels. Oxygen concentration in the atmosphere in a low-pressure weather system (28.5inHg at sea level) is similar to the oxygen levels (97% of normal) found at an elevation of 400m above sea level. Lower blood oxygen concentration (SpO_2_) is associated with lower barometric pressure [29]. An imaging study of mice subjected to low levels of O_2_ found that macro-vasculature decreased in volume, while microvasculature blood flow increased [43]. Tissue requires more blood to deliver the same amount of oxygen in a low-O_2_ environment and may thus cause a small and temporary change in brain volume. However, this does not explain why the cerebellum follows a different pattern from the rest of the brain.

Since brain volumes have been assumed to be static except for the effects of aging, few if any studies have examined temporally fine-grained (daily or weekly) MRI scans for extended periods of time. Replication of the changes found in this analysis would be best tested using a single subject, or set of subjects, scanned daily throughout an entire year. Such data would confirm whether the effects seen in this study are a biological effect or are only found at a group-level in a heterogeneous group. Either finding would be important to the interpretation of large-scale heterogeneous studies.

Investigating the biological cause of such large volume changes may be clinically relevant, including why volume changes are observed in opposite directions in the supratentorium vs infratentorium. Investigating these changes further may be informative for seasonal disorders or discover previously unknown seasonal effects on other diseases. From a purely statistical standpoint, adding season and weather variables to big data analyses may improve accuracy, especially if the analysis includes geographic sites that experience wide variation in these variables.

## Limitations

Many demographic and phenotypic variables were not collected for all subjects used in this analysis. Variables such as race, ethnicity, education, BMI, medication, menstrual cycle, recreational drug use, and smoking status were only collected on a small subset of subjects, and those subsets often did not overlap, were inconsistently recorded between projects, or were mostly just not available because of inaccessibility to paper records.

## Data Availability Statement

The data that support the findings of the study are available on request from the corresponding author (GB). The data are not publicly available due them containing information that could compromise research participant privacy/consent.

## Acknowledgements

We would like to acknowledge the entire past and present staff of the Olin Neuropsychiatry Research Center for 15 years’ worth of MRI data collection and the individual projects under which the data was collected.

## References

1. Hoffmann, J., et al., The influence of weather on migraine - are migraine attacks predictable? Ann Clin Transl Neurol, 2015. 2(1): p. 22–8.

2. Akutagawa, O., H. Nishi, and K. Isaka, Spontaneous delivery is related to barometric pressure. Arch Gynecol Obstet, 2007. 275(4): p. 249–54.

3. Ma, J. and X. Zhang, [The relationship between season/latitude and multiple sclerosis]. Zhonghua Nei Ke Za Zhi, 2015. 54(11): p. 945–8.

4. Spelman, T., et al., Seasonal variation of relapse rate in multiple sclerosis is latitude dependent. Ann Neurol, 2014. 76(6): p. 880–90.

5. Hallam, K.T., et al., Seasonal influences on first-episode admission in affective and non-affective psychosis. Acta Neuropsychiatr, 2006. 18(3-4): p. 154–61.

6. Owens, N. and P.D. McGorry, Seasonality of symptom onset in first-episode schizophrenia. Psychol Med, 2003. 33(1): p. 163–7.

7. Lim, A.S., et al., Diurnal and seasonal molecular rhythms in human neocortex and their relation to Alzheimer’s disease. Nat Commun, 2017. 8: p. 14931.

8. Miller, M.A., et al., Photoperiod is associated with hippocampal volume in a large community sample. Hippocampus, 2015. 25(4): p. 534–43.

9. Nakamura, K., et al., Diurnal fluctuations in brain volume: Statistical analyses of MRI from large populations. Neuroimage, 2015. 118: p. 126–32.

10. Trefler, A., et al., Impact of time-of-day on brain morphometric measures derived from T1-weighted magnetic resonance imaging. Neuroimage, 2016. 133: p. 41–52.

11. Meyer, C., et al., Seasonality in human cognitive brain responses. Proc Natl Acad Sci U S A, 2016. 113(11): p. 3066–71.

12. Lim, A.S.P., et al., Seasonal plasticity of cognition and related biological measures in adults with and without Alzheimer disease: Analysis of multiple cohorts. PLoS Med, 2018. 15(9): p. e1002647.

13. Lazaro, J., et al., Profound seasonal changes in brain size and architecture in the common shrew. Brain Struct Funct, 2018. 223(6): p. 2823–2840.

14. Luo, Y., et al., Seasonality and brain size are negatively associated in frogs: evidence for the expensive brain framework. Sci Rep, 2017. 7(1): p. 16629.

15. Clayton, N.S., J.C. Reboreda, and A. Kacelnik, Seasonal changes of hippocampus volume in parasitic cowbirds. Behav Processes, 1997. 41(3): p. 237–43.

16. Edwards, F.A. and P.W. Gage, Seasonal changes in inhibitory currents in rat hippocampus. Neurosci Lett, 1988. 84(3): p. 266–70.

17. Iaskin, V.A., [Seasonal changes in hippocampus size and spatial behaviour in mammals and birds]. Zh Obshch Biol, 2011. 72(1): p. 27–39.

18. Fan, C., et al., Reversible Brain Abnormalities in People Without Signs of Mountain Sickness During High-Altitude Exposure. Sci Rep, 2016. 6: p. 33596.

19. Van Ombergen, A., et al., Brain Tissue-Volume Changes in Cosmonauts. N Engl J Med, 2018. 379(17): p. 1678–1680.

20. Fischl, B., FreeSurfer. Neuroimage, 2012. 62(2): p. 774–81.

21. Book, G.A., et al., Neuroinformatics Database (NiDB)--a modular, portable database for the storage, analysis, and sharing of neuroimaging data. Neuroinformatics, 2013. 11(4): p. 495–505.

22. Fischl, B., et al., Whole brain segmentation: automated labeling of neuroanatomical structures in the human brain. Neuron, 2002. 33(3): p. 341–55.

23. Reuter, M., et al., Head motion during MRI acquisition reduces gray matter volume and thickness estimates. Neuroimage, 2015. 107: p. 107–115.

24. Jenkinson, M., et al., Improved optimization for the robust and accurate linear registration and motion correction of brain images. Neuroimage, 2002. 17(2): p. 825–41.

25. M. C. Peel, B.L.F., T. A. McMahon, Updated world map of the Koppen-Geiger climate classification. Hydrology and Earth Science Systems, 2007. 11: p. 1633–1644.

26. Meyers, S.M., et al., Does hydration status affect MRI measures of brain volume or water content? J Magn Reson Imaging, 2016. 44(2): p. 296–304.

27. Streitburger, D.P., et al., Investigating structural brain changes of dehydration using voxel-based morphometry. PLoS One, 2012. 7(8): p. e44195.

28. Fan, J., F. Han, and H. Liu, Challenges of Big Data Analysis. Natl Sci Rev, 2014. 1(2): p. 293–314.

29. Pope, C.A.r., et al., Oxygen saturation, pulse rate, and particulate air pollution: A daily time-series panel study. Am J Respir Crit Care Med, 1999. 159(2): p. 365–72.

30. Jehn, M., et al., The effect of ambient temperature and barometric pressure on ambulatory blood pressure variability. Am J Hypertens, 2002. 15(11): p. 941–5.

31. van Erp, T.G., et al., Subcortical brain volume abnormalities in 2028 individuals with schizophrenia and 2540 healthy controls via the ENIGMA consortium. Mol Psychiatry, 2016. 21(4): p. 547–53.

32. Schmaal, L., et al., Subcortical brain alterations in major depressive disorder: findings from the ENIGMA Major Depressive Disorder working group. Mol Psychiatry, 2016. 21(6): p. 806–12.

33. Whelan, C.D., et al., Structural brain abnormalities in the common epilepsies assessed in a worldwide ENIGMA study. Brain, 2018. 141(2): p. 391–408.

34. Mackey, S., et al., Mega-Analysis of Gray Matter Volume in Substance Dependence: General and Substance-Specific Regional Effects. Am J Psychiatry, 2018: p. appiajp201817040415.

35. Kurlansik, S.L. and A.D. Ibay, Seasonal affective disorder. Am Fam Physician, 2012. 86(11): p. 1037–41.

36. Ozeki, K., et al., Weather and headache onset: a large-scale study of headache medicine purchases. Int J Biometeorol, 2015. 59(4): p. 447–51.

37. Cioffi, I., et al., Effect of weather on temporal pain patterns in patients with temporomandibular disorders and migraine. J Oral Rehabil, 2017. 44(5): p. 333–339.

38. Becker, W.J., Weather and migraine: can so many patients be wrong? Cephalalgia, 2011. 31(4): p. 387–90.

39. Annweiler, C., et al., Vitamin D-related changes in intracranial volume in older adults: a quantitative neuroimaging study. Maturitas, 2015. 80(3): p. 312–7.

40. Klingberg, E., et al., Seasonal variations in serum 25-hydroxy vitamin D levels in a Swedish cohort. Endocrine, 2015. 49(3): p. 800–8.

41. Kristal-Boneh, E., et al., Summer-winter variation in 24 h ambulatory blood pressure. Blood Press Monit, 1996. 1(2): p. 87–94.

42. Stergiou, G.S., et al., Seasonal variation in meteorological parameters and office, ambulatory and home blood pressure: predicting factors and clinical implications. Hypertens Res, 2015. 38(12): p. 869–75.

43. Jia, Y., et al., Responses of peripheral blood flow to acute hypoxia and hyperoxia as measured by optical microangiography. PLoS One, 2011. 6(10): p. e26802.

